# Memory dysfunction and psychiatric outcomes in anti-NMDA receptor encephalitis are linked to altered structural brain complexity

**DOI:** 10.1101/2025.09.22.677264

**Authors:** Stephan Krohn, Guido Cammà, Wei Zhao, Amy Romanello, Katharina Wurdack, Ole Jonas Böken, Sophia Rekers, Friedemann Paul, Harald Prüß, Christopher R. Madan, Carsten Finke

**Author notes:** Corresponding author: Carsten Finke Department of Neurology Charité-Universitätsmedizin Berlin Charitéplatz 1 10117 Berlin Germany.

## Abstract

**Introduction:** Most patients with anti-N-methyl-D-aspartate receptor encephalitis (NMDARE) experience long-term neuropsychiatric sequelae despite immunotherapy. However, how these residual symptoms relate to structural brain changes in the post-acute phase remains unclear. Recently, fractal dimensionality (FD) has emerged as a sensitive imaging marker of structural brain complexity in related conditions but has not been explored in autoimmune encephalitis.

**Methods:** This cross-sectional study combined clinical, cognitive, and neuroimaging analyses in 70 patients with post-acute NMDARE (median time from onset: 22 months), and 70 healthy controls matched for age (*t*=-0.57, *p*=0.57) and sex (*χ*^2^=0, *p*=1). High-resolution T1-weighted magnetic resonance imaging (MRI) data were analyzed with FreeSurfer and computational fractal analysis. Clinical outcomes were assessed with the modified Rankin Scale (mRS) and Clinical Assessment Scale in Autoimmune Encephalitis (CASE). Psychiatric manifestations underwent phenotypical analyses, and memory performance was assessed with standardized neuropsychological tests.

**Results:** Patients with NMDARE were severely affected at peak illness but showed substantial overall improvement in the post-acute stage (median mRS at peak=5, post-acute=1, *z*=-7.2, *p*<0.001; median CASE at peak=11, post-acute=1, *z*=-7.2, *p*<0.001). However, 67% of patients showed residual CASE symptoms, with memory dysfunction (61%) and psychiatric symptoms (36%) representing the most prevalent domains. Therein, psychiatric symptoms showed a phenotypical shift from a ‘schizophrenia-like’ phenotype at peak illness to an affective phenotype in the post-acute stage. Neuroimaging uncovered a characteristic pattern of reduced structural brain complexity, including the hippocampus bilaterally and a fronto-cingulo-temporal cluster in cortical gray matter and cerebral white matter. Importantly, normative modeling revealed that patients with residual symptoms showed stronger alterations of brain complexity than those without (psychiatric: *t*=-2.65, *p*=0.010; memory: *t*=-3.98, *p*<0.001; both vs. no symptoms: *t*=-5.46, *p*<0.001). Similarly, patients with stronger alterations of brain complexity showed lower scores of visuospatial and verbal memory (all *p_FDR_*<0.015).

**Discussion:** Post-acute NMDARE is characterized by systematic reductions in structural brain complexity, consistently involving previously implicated regions while identifying changes in the cingulate cortex as a new morphological correlate. These changes are linked to residual psychiatric symptoms and memory dysfunction, highlighting FD as a promising new imaging marker of long-term outcomes. Our findings suggest that current treatment strategies may be insufficient to fully address the residual symptom burden after the acute phase of NMDARE.

## Introduction

Anti-N-methyl-D-aspartate receptor encephalitis (NMDARE) represents the most common form of autoimmune encephalitis (AE)^1^. First discovered in 2007^2^, NMDARE is caused by autoantibodies against the GluN1 subunit of the NMDA receptor^3^, which plays a key role in glutamatergic signaling, long-term potentiation, and synaptic plasticity across the brain^4,5^. Clinically, NMDARE is characterized by a severe neuropsychiatric syndrome including cognitive dysfunction, psychiatric symptoms, seizures, autonomic dysregulation, and decreased levels of consciousness, which frequently necessitates intensive care^1,6,7^. Although significant advances have been made in managing the acute disease phase^7,8^, our understanding of long-term outcomes remains incomplete.

Recent evidence suggests that, although most patients achieve favorable neurological outcomes overall, long-term recovery is often protracted, and many patients experience residual symptoms for years after disease onset, despite immunotherapy^9,10^. These residual symptoms include persistent memory impairment, psychiatric symptoms as well as difficulties in resuming premorbid levels of psychosocial function, occupational activities, and overall quality of life^9–13^.

Despite this clinical relevance, it remains largely unclear how residual symptoms in NMDARE relate to long-term changes in brain structure. Notably, conventional clinical magnetic resonance imaging (MRI) is frequently normal or unspecific in the acute disease phase^14,15^, which has motivated advanced neuroimaging studies to characterize disease-related brain changes beyond overt lesions. In this context, structural complexity analysis has recently emerged as a powerful new tool to describe brain morphology from anatomical MRI data^16,17^. Specifically, fractal dimensionality (FD) represents a marker of structural complexity which yields a quantitative account of brain shape and has shown high sensitivity in capturing age-related changes of brain morphology across the lifespan^17–20^ and in detecting structural brain changes across a variety of clinical conditions, including psychiatric and neuroimmunological disorders^21,22^. Despite these promising reports in related conditions, FD has remained unexplored in AE.

Against this background, the aims of our study were threefold: (i) to analyze structural brain complexity in a large sample of healthy individuals and patients with NMDARE; (ii) to characterize patient outcomes from peak illness to the post-acute visits at which the MRI was performed; and (iii) to study how structural brain changes are related to residual symptoms in post-acute NMDARE.

## Methods

### Study population

We prospectively included 70 patients with NMDARE referred to our reference center at the Department of Neurology, Charité-Universitätsmedizin Berlin, Germany. Data were acquired between July 2011 and March 2021. All patients fulfilled current diagnostic criteria^23^ and tested positive for anti-NMDA receptor antibodies in CSF. All patients were in the post-acute disease stage, defined as having been discharged from acute hospitalization for at least 1 month. Study visits occurred at a median of 22.2 months since disease onset (IQR: 8.5-42.5; 87% of patients >6 months). Patients were matched to 70 healthy control (HC) participants for sex and age using logistic regression propensity scores and nearest-neighbor matching, as implemented in the MatchIt package (version 4.5.5) for R (version 4.4.0). Therein, exact matching was applied for participants’ sex-at-birth, resulting in identical sex ratios in both groups. Study approval was obtained by the institutional ethics committee (EA1/206/10, EA1/095/12, EA1/105/16, EA4/011/19), and all participants gave written informed consent.

### Clinical scores

Clinical data were collected using standardized case report forms. The modified Rankin Scale (mRS) and the Clinical Assessment Scale in Autoimmune Encephalitis^11^ (CASE) were scored retrospectively by two investigators (GC, KW). Clinical data for mRS scores were available for 69/70 patients at peak illness, and for all patients at post-acute visit. Similarly, data for CASE scores were available for 68/70 patients at peak illness, and for all patients at post-acute follow-up.

Given the high prevalence of psychiatric CASE symptoms, we classified these symptoms at a higher clinical granularity. Two experienced researchers (KW, OJB) independently evaluated the clinical information for the presence of the following symptom categories: psychosis, psychomotor symptoms, catatonia, dissociative symptoms, personality changes, manic symptoms, depressivity, affective dysregulation, suicidality/self-harm, anxiety-related symptoms, aggressivity, and impaired impulse control. Clinical data allowed for a further subdivision of psychosis (hallucinations, delusions, paranoia, other) and psychomotor symptoms (increase vs. decrease). Symptoms that did not fit any of the categories were rare, unspecific (predominantly confusion), and omitted from quantitative analyses. The independent evaluations at each timepoint (i.e., acute and post-acute) were cross-checked by the respective other researcher and subsequently reviewed by two initially blinded researchers (SK, CF).

### Memory performance

Patients’ memory performance at the time of MRI was assessed with standardized neuropsychological examination. Specifically, visuospatial memory was evaluated with the Rey-Osterrieth Complex Figure test (ROCF; immediate recall and delayed recall), and verbal episodic memory was assessed with the German edition of the Rey Auditory Verbal Learning Test (RAVLT; sum score, post-interference trial, delayed recall). Besides analyzing these raw values, we conducted a normative interpretation of memory performance based on the respective delayed recall scores. To this end, individual scores were converted to percentile ranks with age-specific norms and labeled as ‘low performance’ at values <25th percentile and ‘high performance’ at ≥75th percentile, in line with recent neuropsychological consensus guidelines^24^.

### Neuroimaging analyses

Anatomical MRI data were acquired with a 3T Trim Trio scanner (Siemens, Erlangen, Germany) at the Berlin Center for Advanced Neuroimaging. High-resolution T1-weighted images were obtained using a magnetization-prepared rapid gradient echo sequence (TR/TE/TI=1900/2.55/900ms, FOV=240×240mm^2^, matrix size=240×240, 176 slices, slice thickness=1mm). These structural images were processed with comprehensive FreeSurfer reconstruction (version 6.0.0)^25^, including volumetric white matter (WM) parcellation and hippocampal subfield segmentation. Regions of interest (ROI) were based on the Desikan-Killiany atlas^26^, and global parcellations of cortical gray matter (GM) and cerebral WM were derived using the respective voxel index supersets^27^.

The resulting volumetric segmentations were subsequently passed to region-wise structural complexity analysis. Specifically, we estimated the FD of each ROI from the spatial scaling properties of the corresponding segmentation mask^16,17^, using the dilation algorithm for filled structures from the openly available *calcFD* toolbox (https://github.com/cMadan/calcFD) for MATLAB (The MathWorks Inc., Natick, Massachusetts, USA). Briefly, this algorithm implements an iterative convolution of the segmentation mask with a set of spatial kernels to estimate the power-law relationship between the size of the spatial scale and the count of scaled measurement units^18^. For fractal analysis of structural MRI, these spatial scales correspond to a range of voxel sizes, which can be expressed as 2^*k*^, *k*∈ℕ_0_. Here we apply the default range of *k* = 0, 1,…, 4, as previously validated^17–20,28^. The FD estimate is then computed as the slope of the relationship between scale size and unit count in log-log space^16–18^. This procedure thus yields a single FD value as a measure of the region’s structural complexity, with higher values reflecting more irregular and space-filling shapes^17^.

### Normative modelling

To quantify global changes of structural brain complexity, we applied a normative modelling framework that we recently developed to compare the whole-brain FD profiles of individual participants to those of a specified reference population^17^. For clinical neuroimaging studies, this reference population is defined as the HC group^29^, and normative FD values are computed as the median over these healthy participants. This approach thus yields a reference vector of FD values over all brain regions, and patients’ overall change in structural brain complexity is quantified as the rank correlation distance between their individual FD values and this normative reference vector. Consequently, we obtain a single deviation index for each patient, which quantifies how much their FD values across the whole brain deviate from the healthy reference. Notably, this approach has shown high sensitivity in a recent neurodevelopmental study^17^ and a related application in anti-LGI1 encephalitis^29^ and features several theoretical advantages: First, the index considers all regions simultaneously, which reduces the high dimensionality of brain space and avoids multiple ROI-wise testing. Second, it allows for individual deviation patterns as the approach is agnostic to which particular regions drive the deviation in a given patient. Third, increases and decreases jointly contribute to the deviation index, such that it explicitly captures bidirectional deviations that can remain undetected in group-level analyses.

### Region-to-network mapping

To investigate how structural brain changes relate to functional systems, we applied a recent region-to-network mapping procedure^29^. Briefly, this procedure maps anatomical GM regions to one of the seven resting-state networks in the Multiresolution Intrinsic Segmentation Template^30^ which includes the cerebellum, mesolimbic network, somatomotor network, visual network, default mode network, frontoparietal/visual-downstream network, and ventral attention network/salience network/basal ganglia/thalamus as functional clusters. For further details, please refer to Krohn, Müller-Jensen et. al. (2025)^29^.

### Statistical analysis

Continuous relationships were assessed with product-moment correlations, frequencies with *χ*^2^ proportion tests, and univariate between-group comparisons with independent-sample *t*-tests and standardized effect sizes (Cohen’s *d*). Clinical scores (mRS, CASE) assessed once at peak illness and once at post-acute follow-up were compared with signed rank tests. Group differences in structural complexity estimates were assessed with a non-parametric permutation test. The null hypothesis under this approach posits that there is no group difference and, consequently, it is irrelevant whether a particular participant is labeled a patient or control. To test this hypothesis, one estimates the ensuing null distribution by randomly permutating the group labels and recalculating the statistic for every permutation. The *p*-value of this test is then given by the number of instances in which the randomly permuted statistics surpass the value of the empirically observed statistic, divided by the total number of random permutations (here n=5000). The standardized effect size of these comparisons was computed as 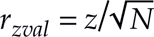, where *z* corresponds to the empirical test statistic and *N* = *n*_*HC*_+ *n*_*NMDARE*_ to the total sample size. Moreover, we calculated the 95% confidence interval of these estimates with the adjusted bootstrap percentile method and 1000 bootstrap iterations, using the rstatix package (version 0.7.2). When a variable was repeatedly tested against multiple other measures, we applied the Benjamini-Hochberg procedure to control the false discovery rate (FDR).^31^

## Code availability

Resources supporting the findings of this study will be made available on the Open Science Framework upon acceptance.

## Results

### Patient characteristics

The study population included 70 patients with NMDARE and 70 healthy control participants (HC), matched for age (*t*=-0.57, *p*=0.57) and sex (*χ*^2^=0, *p*=1). In line with previous reports^6^, most patients were female (86%) and in early adulthood at disease onset (mean 25.7±10.1 years). Patients were severely affected in the acute disease phase (median mRS: 5; median CASE: 11). The most common clinical manifestations included psychiatric symptoms (97%), cognitive deficits (94%), specifically memory dysfunction (91%), and seizures (78%). Treatment included first-line (97%), second-line (58%), and third-line (4%) immunotherapy as well as anticonvulsive and antipsychotic medication (75% each). Further clinical characteristics are summarized in **Table 1**.

**Table 1.** Clinical characteristics of the patient sample.

| Characteristic | Patients, no. (%) |
| --- | --- |
|  | NMDARE (n = 70) |
| **Demographics** |  |
| Age at onset, years (mean ± SD; range) | 25.7 ± 10.1 (14-67) |
| Age at MRI, years (mean ± SD; range) | 28.1 ± 10.2 (15-67) |
| Sex (female / male) (%) | 60 (86) / 10 (14) |
| **Clinical presentation in acute stage** |  |
| Prodromal symptoms (%) | 33 / 66 (50) |
| Memory dysfunction (%) | 63 / 69 (91) |
| Psychiatric symptoms (%) | 67 / 69 (97) |
| Seizures (%) | 53 / 68 (78) |
| Sleep disturbance (%) | 37 / 59 (63) |
| Movement disorders (%) | 35 / 66 (53) |
| Autonomic dysfunction (%) | 32 / 69 (46) |
| Impaired higher-order cognition (%) | 62 / 66 (94) |
| **Treatment** |  |
| Days from onset to treatment, median (IQR) | 31 (14-164) |
| **First-line therapy** (%) | 66 / 68 (97) |
| IV steroids (%) | 44 / 68 (65) |
| Plasmapheresis (%) | 33 / 68 (49) |
| Immunoadsorption (%) | 12 / 68 (18) |
| IV immunoglobulins (%) | 29 / 68 (43) |
| **Second-line therapy** (%) | 39 / 67 (58) |
| Rituximab (%) | 27 / 67 (40) |
| Cyclophosphamide (%) | 8 / 67 (12) |
| Sustained oral steroids (%) | 6 / 67 (9) |
| Azathioprine (%) | 5 / 67 (7) |
| Methotrexate (%) | 3 / 67 (4) |
| **Third-line therapy** (%) | 3 / 67 (4) |
| Bortezomib (%) | 3 / 67 (4) |
| Anticonvulsive medication (%) | 50 / 67 (75) |
| Antidepressant medication (%) | 22 / 66 (33) |
| Antipsychotic medication (%) | 49 / 65 (75) |
| **Months from onset to MRI**, median (IQR) | 22 (8.5-42.5) |
| **Outcome** |  |
| mRS score at peak, median (range) | 5 (2-5) |
| mRS score at MRI, median (range) | 1 (0-4) |
| CASE sum score at peak, median (range) | 11 (1-27) |
| CASE sum score at MRI, median (range) | 1 (0-12) |
| Relapse (%) | 9 / 68 (13) |

Abbreviations: CASE, Clinical Assessment Scale for Autoimmune Encephalitis; IQR, interquartile range; IV, intravenous; MRI, magnetic resonance imaging; mRS, modified Rankin Scale; NMDARE, anti-N-methyl-D-aspartate receptor encephalitis; SD, standard deviation.

### Structural complexity analysis

We first investigated whether patients with NMDARE exhibit systematic alterations of structural brain complexity. As summarized in **Figure 1**, we observed a characteristic pattern of brain-wide FD reductions in patients compared to HC. Specifically, patients showed significantly lower FD across the cerebral cortex (**Fig. 1A**, *left*), and these reductions clustered bilaterally in the frontal lobe (strongest effect left: caudal middle frontal gyrus: *z*=-2.48, *p_perm_*=0.006; right: rostral middle frontal gyrus: *z*=-2.85, *p_perm_*=0.002), the temporal lobe (left: entorhinal cortex: *z*=-3.17, *p_perm_*=0.001; right: entorhinal cortex: *z*=-2.71, *p_perm_*=0.004), and the cingulate (left: isthmus: *z*=-3.28, *p_perm_*<0.001; right: posterior cingulate: *z*=-2.43, *p_perm_*=0.008). Notably, this fronto-cingulo-temporal cluster of FD reductions was corroborated in global parcellations of cortical GM (frontal: *z*=-2.21, *p_perm_*=0.011; temporal: *z*=-2.22, *p_perm_*=0.013; cingulate: *z*=-3.22, *p_perm_*<0.001; **Fig. 1A**, *right*).

**Figure 1.**
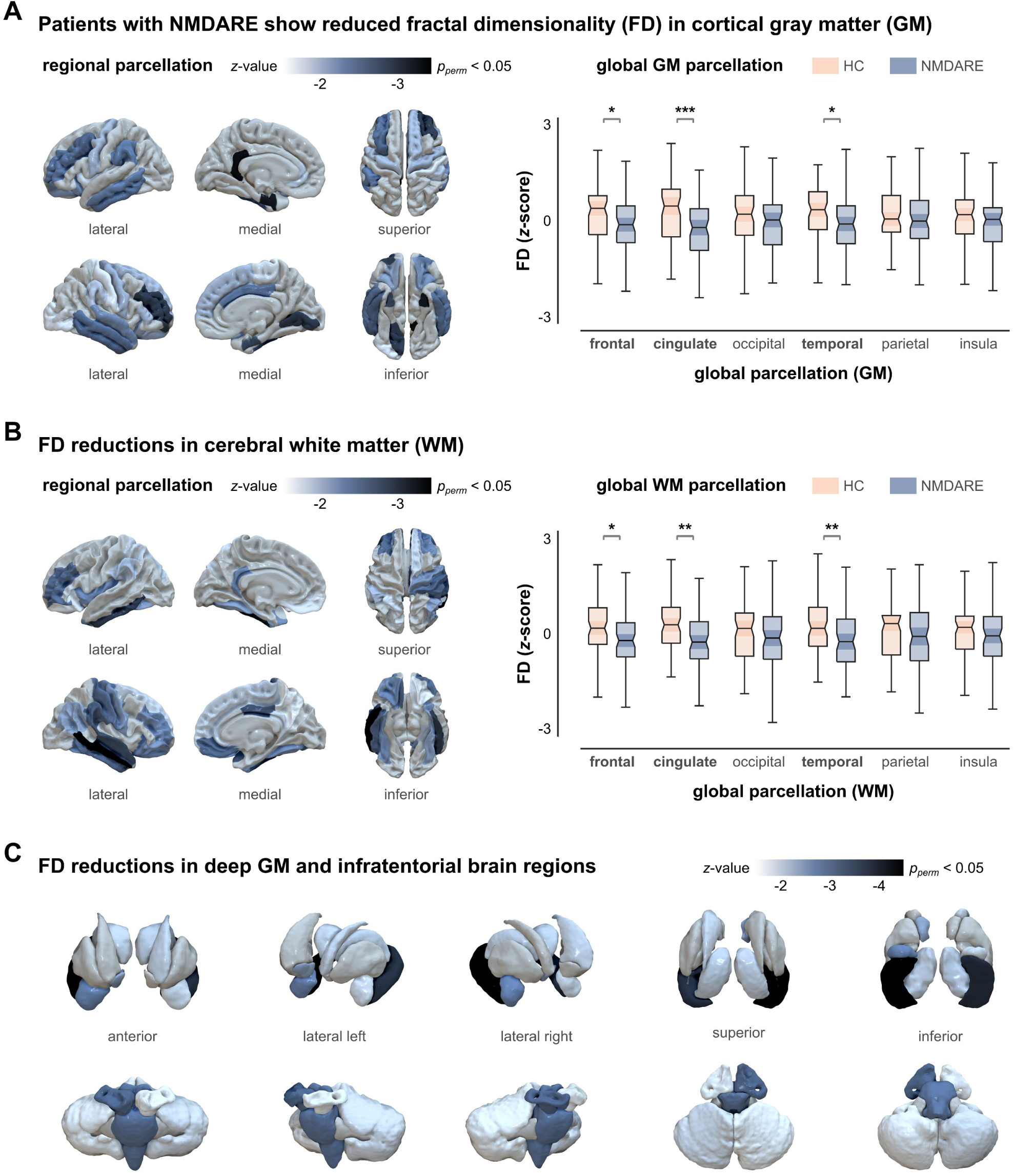
Patients with NMDARE show systematic reductions of structural brain complexity. **(A)**, Comparing fractal dimensionality (FD) values of cortical gray matter (GM) between patients with anti-NMDA receptor encephalitis (NMDARE) and healthy control participants (HC; n=70 each; threshold: permuted *p*<0.05). The left panel corresponds to regional GM parcellation based on the Desikan-Kiliany atlas^26^, and the right panel to global cortex parcellation. **(B)**, Group differences in cerebral white matter (WM). **(C)**, Group differences in deep GM and infratentorial regions.

A remarkably similar pattern emerged in cerebral WM (**Fig. 1B**, *left*), where FD reductions again clustered in the frontal lobe (left: rostral middle frontal: *z*=-2.43, *p_perm_*=0.006; right: medial orbitofrontal: *z*=-2.51, *p_perm_*=0.004), temporal lobe (left: inferior temporal: *z*=-2.88, *p_perm_*=0.002; right: middle temporal: *z*=-3.72, *p_perm_*<0.001), and cingulate (left: isthmus: *z*=-2.22, *p_perm_*=0.014; right: posterior cingulate: *z*=-2.68, *p_perm_*=0.003). Global parcellations of cerebral WM again confirmed this spatial pattern of FD reductions (frontal: *z*=-2.06, *p_perm_*=0.020; temporal: *z*=-2.34, *p_perm_*=0.007; cingulate: *z*=-2.64, *p_perm_*=0.003; **Fig. 1B**, *right*). Additionally, patients showed reduced FD in deep GM areas and infratentorial regions (**Fig. 1C**), where the most pronounced FD reductions across the whole brain were observed in the hippocampus bilaterally (left: *z*=-3.82, *p_perm_*<0.001; right: *z*=-4.54, *p_perm_*<0.001).

Given these strong hippocampal effects, we next examined whether the observed FD reductions were uniformly distributed across the whole hippocampus or varied by hippocampal subfield. As shown in **Figure 2**, we observed a subfield-specific magnitude of hippocampal FD reductions: While some regions showed little-to-no difference between patients and controls (e.g., subiculum, parasubiculum), the strongest FD reductions were observed in the CA1 subregion, dentate gyrus, and molecular layer.

**Figure 2.**
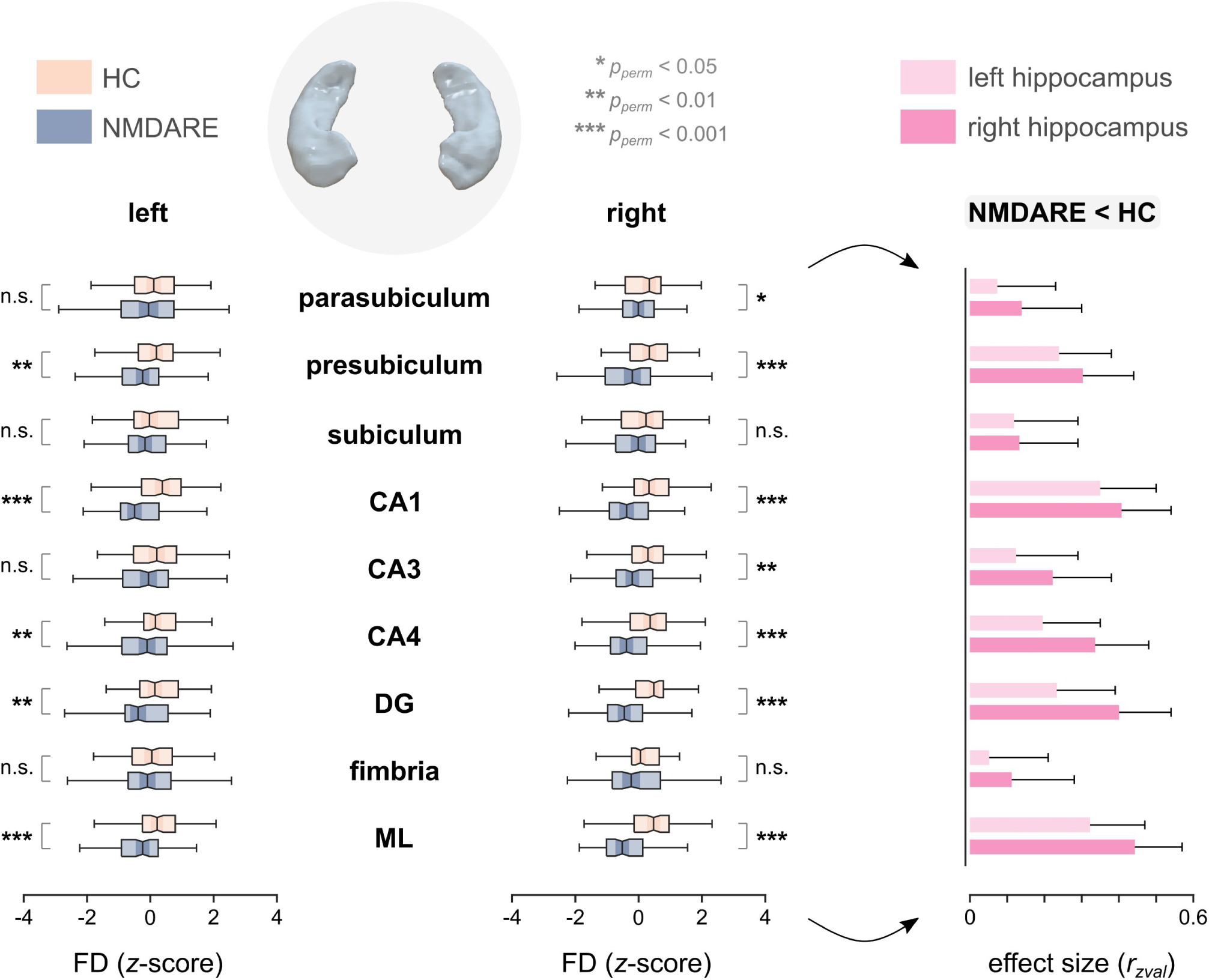
The magnitude of hippocampal FD reductions varies across subfields. Comparing fractal dimensionality (FD) values of patients with anti-NMDA receptor encephalitis (NMDARE) and healthy control participants (HC; n=70 each) across hippocampal subfields. The right panel shows the standardized effect sizes of subfield-specific FD reductions in NMDARE. Error bars correspond to the upper bound of the bootstrapped 95% confidence interval. Abbreviations: CA, cornu ammonis; DG, dentate gyrus; ML, molecular layer.

In sum, these analyses revealed a characteristic anatomical pattern of FD reductions, including the hippocampus bilaterally and a fronto-cingulo-temporal cluster in the cerebrum. Therefore, we next studied how these FD reductions are distributed across functional brain systems. To this end, we applied a recent region-to-network mapping procedure, which assigns structural changes in GM regions to one of seven functional clusters^29^.

As shown in **Figure 3**, the most prominent cluster of FD reductions corresponds to the mesolimbic network, which included both the highest proportion of affected regions and the most pronounced effects in between-group testing (left and right hippocampus; **Fig. 3**, *top*). Notably, both the ventral attention / salience cluster (second in proportion) and the default mode network (third) showed their respective strongest effects in the cingulate cortex (**Fig. 3**, *center*), while frontal FD reductions belonged predominantly to the frontoparietal network (**Fig. 3**, *bottom*). Overall, FD reductions thus clustered in several higher-order functional systems, while primary networks (visual, somatomotor) were comparatively less affected.

**Figure 3.**
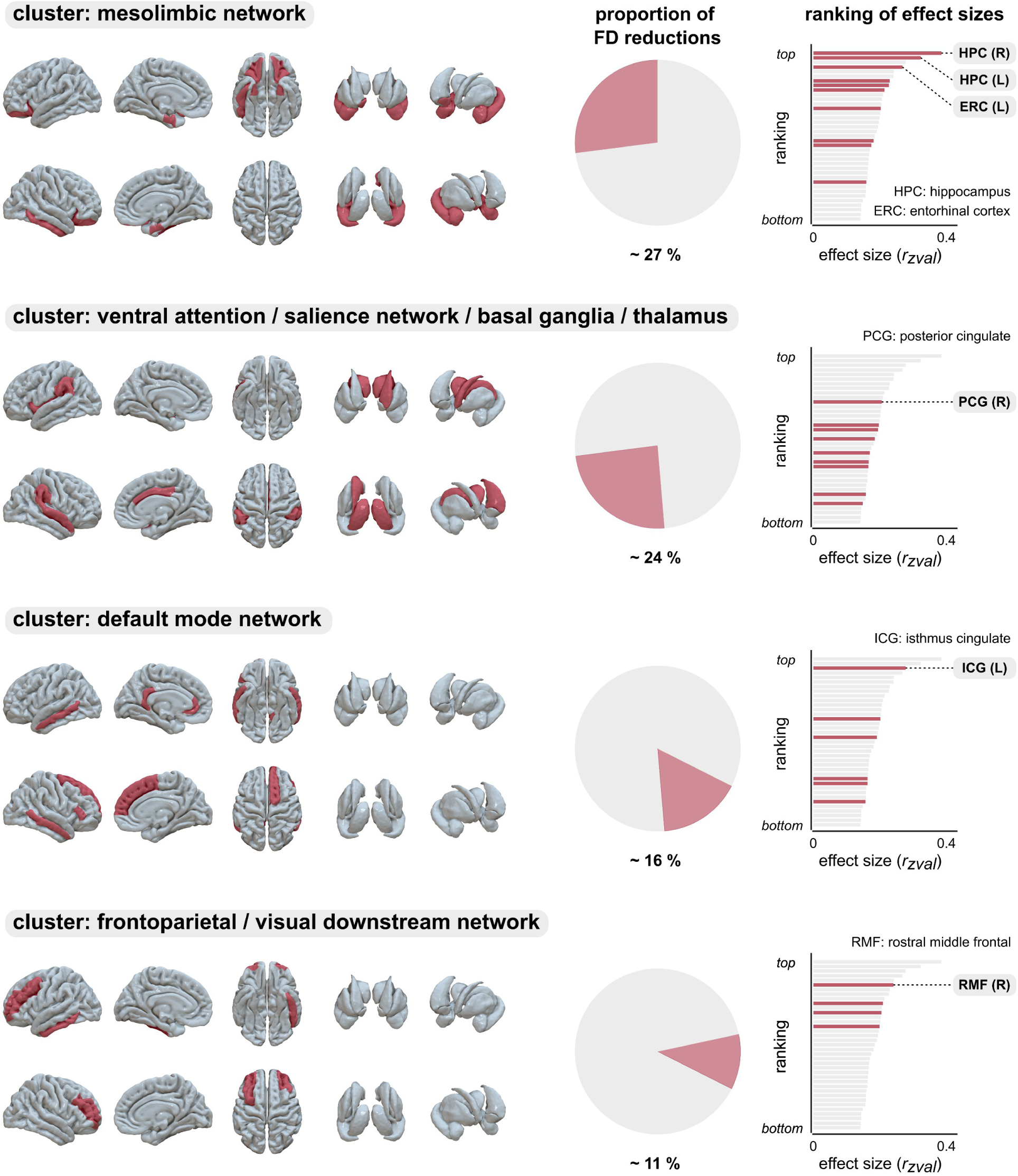
FD reductions in gray matter regions cluster over functional systems. The brain plots visualize the network assignment of gray matter (GM) regions with significant between-group differences (see Fig. 1). Functional assignments rest on a recent region-to-network mapping procedure^29^ and are ordered by the proportion of affected regions in the cluster over all affected regions (middle panels). The right panels show the relative effect ranking of significant FD reductions, where regions belonging to the cluster are highlighted and regions with the respective strongest effects are labelled.

### Patient outcomes and residual symptoms

Next, we studied how these changes in structural brain complexity relate to patient outcomes. As detailed in Table 1, patients were severely affected at peak illness (median mRS=5) but showed significant clinical improvement at the post-acute study visits when MRI data were acquired (median mRS=1; peak illness vs. post-acute: *z*=-7.2, *p*<0.001, n=69).

This finding was corroborated by the more disease-specific CASE score, summarized in **Figure 4**. Overall, the CASE sum score decreased markedly from peak illness to post-acute follow-up (median peak=11, median post-acute=1; *z*=-7.2, *p*<0.001, n=68; **Fig. 4A**, *left*). However, there was considerable variance in post-acute CASE scores: while 23/70 (33%) patients scored zero on all CASE domains, 47/70 (67%) patients showed a varying degree of residual symptoms at the time of MRI, which was not explained by age (*r*=0.17, *p*=0.16) or time since onset (*r*=-0.02, *p*=0.84).

**Figure 4.**
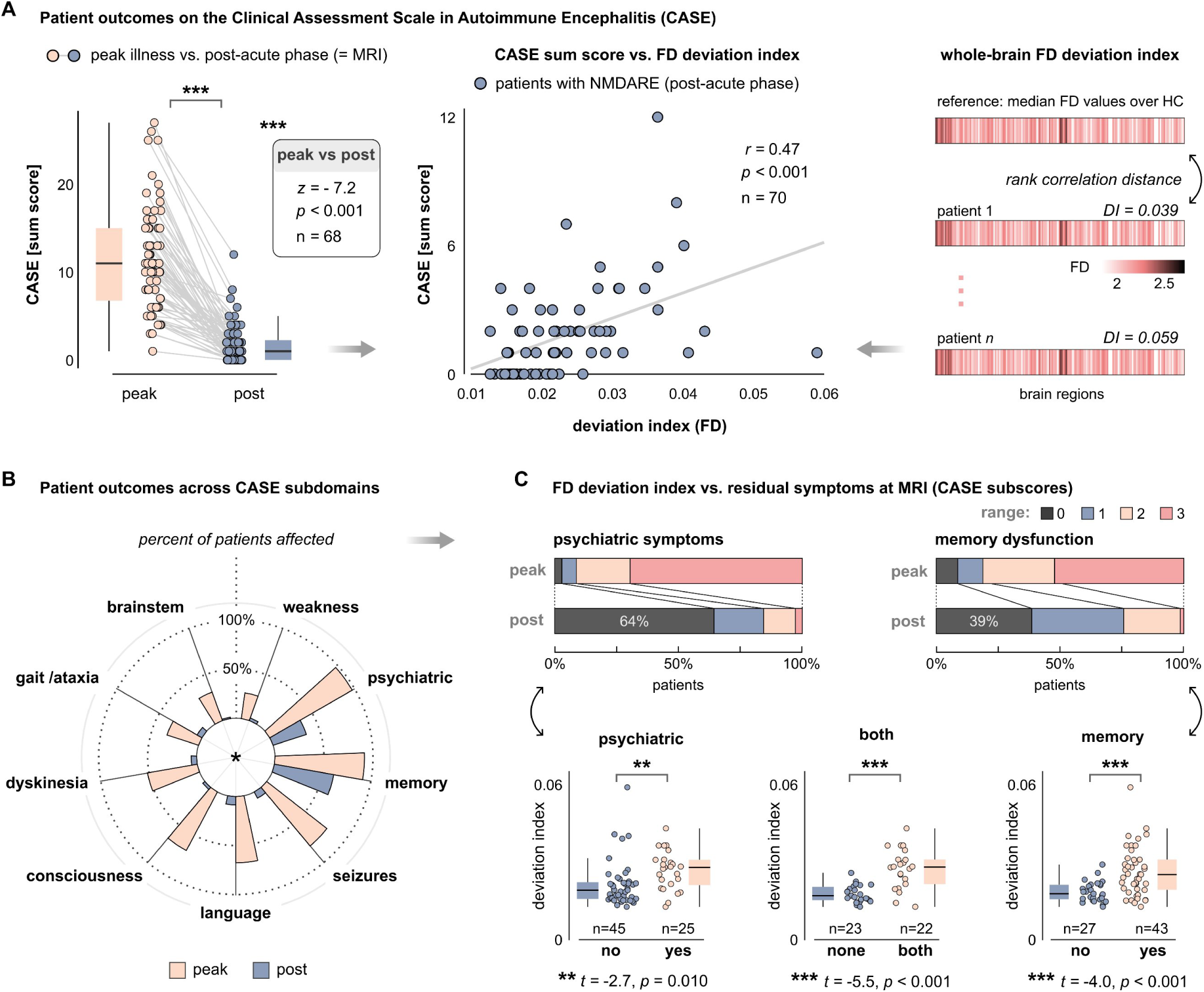
Patients with residual psychiatric and memory symptoms show stronger alterations of structural brain complexity. **(A)**, Patient outcomes on the Clinical Assessment Scale in Autoimmune Encephalitis (CASE). The left panel shows patients’ CASE sum score (possible range: 0-27) from peak illness to the post-acute phase. MRI recordings underlying the imaging analyses were obtained at post-acute follow-up. Peak vs. post-acute scores were compared with a Wilcoxon signed rank test. The right panel illustrates the derivation of the whole-brain fractal dimensionality (FD) deviation index (DI), compared to the reference population of healthy control (HC) participants^17^. The middle panel displays the association between this FD deviation index and post-acute CASE sum scores. **(B)**, Patient outcomes across the CASE subdomains. The polar plot shows the percentage of affected patients at peak illness and post-acute follow-up. **(C)**, The bar plots relate patient outcomes on the two most prevalent clinical subdomains: psychiatric symptoms and memory dysfunction, each scored from 0 (none) to 3 (most severe)^32^. The lower panels show subgroup comparisons of patients who had residual psychiatric symptoms at post-acute follow-up vs. those who did not (left), patients who had residual memory dysfunction vs. those who did not (right), and patients who exhibited both residual psychiatric and memory symptoms vs. those who did not show any CASE symptoms at all (i.e., CASE sum score = 0; middle).

Against this background, we hypothesized that the extent of these residual symptoms would be related to more pronounced changes of structural brain complexity. To test this idea, we applied a recently developed normative modelling framework (see Methods) which compares the whole-brain FD profiles of individual patients to the reference population of HC, yielding an FD deviation index that quantifies patients’ overall change in structural brain complexity (**Fig. 4A**, *right*). We indeed observed a significant positive association between this deviation index and post-acute CASE sum scores (*r*=0.47, *p*<0.001; **Fig. 4A**, *middle*), such that patients with a higher residual symptom burden showed stronger alterations of structural brain complexity.

Therefore, we next studied which of the CASE subdomains drive these residual symptoms (**Fig. 4B**). Notably, patients were affected across all subdomains at peak illness, with psychiatric symptoms (97%), memory dysfunction (91%), and seizures (78%) representing the most prevalent clinical domains. In contrast, residual symptoms at the time of MRI were largely confined to memory dysfunction (61%) and psychiatric symptoms (36%), while all other CASE domains had decreased substantially (<10%).

Accordingly, we focused on these two leading domains specifically (**Fig. 4C**, *top*) and stratified patients into subgroups based on whether they experienced residual symptoms at MRI or not. This subgroup analysis showed significantly stronger alterations of structural brain complexity in patients with residual psychiatric symptoms compared to those without (*t*=-2.65, *p*=0.010, *d*=-0.66; **Fig. 4C**, *bottom-left*). Similarly, we observed stronger complexity alterations in patients with residual memory dysfunction compared to those without (*t*=-3.98, *p*<0.001, *d*=-0.98; **Fig. 4C**, *bottom-right*). Notably, the strongest alterations of structural brain complexity were observed in patients with *both* memory dysfunction and psychiatric symptoms compared to those without *any* residual CASE symptoms (*t*=-5.46, *p*<0.001, *d*=-1.63; **Fig. 4C**, *bottom*-*center*).

Notably, these effects could not be explained by age differences among the patient subgroups (psychiatric yes vs. no: *t*=0.26, *p*=0.79; memory yes vs. no: *t*=-0.63, *p*=0.53; both memory and psychiatric symptoms vs. CASE=0: *t*=-0.26, *p*=0.79). Similarly, we observed no significant sex differences among the subgroups (psychiatric yes vs. no: χ^2^=0.09, *p*=0.76; memory yes vs. no: *t*=1.70, *p*=0.19), although there was a trend towards a higher proportion of males in patients with both memory and psychiatric symptoms compared to those without any residual CASE symptoms (4/18 [18.2%] vs. 1/23 [4.4%], χ^2^=2.18, *p*=0.14).

### Clinical evolution of psychiatric symptoms

Given the link between structural brain complexity and the psychiatric CASE domain, we next sought to characterize psychiatric symptoms in detail and study their temporal evolution from peak illness to the post-acute stage. To this end, we analyzed patients’ psychiatric symptoms across twelve clinical domains, summarized in **Figure 5**. First, we estimated the absolute prevalence of these symptoms across all patients and observed that psychosis was the leading symptom at peak illness (66%), followed by psychomotor symptoms (51%), aggressivity (37%), depressivity (34%) as well as anxiety and personality changes (32% each; **Fig. 5A**, *left*). In contrast, the leading residual symptom in the post-acute stage was affective dysregulation (16%), which predominantly included emotional lability (“mood swings”) and to a lesser extent affective flattening, followed by depressivity (12%).

**Figure 5.**
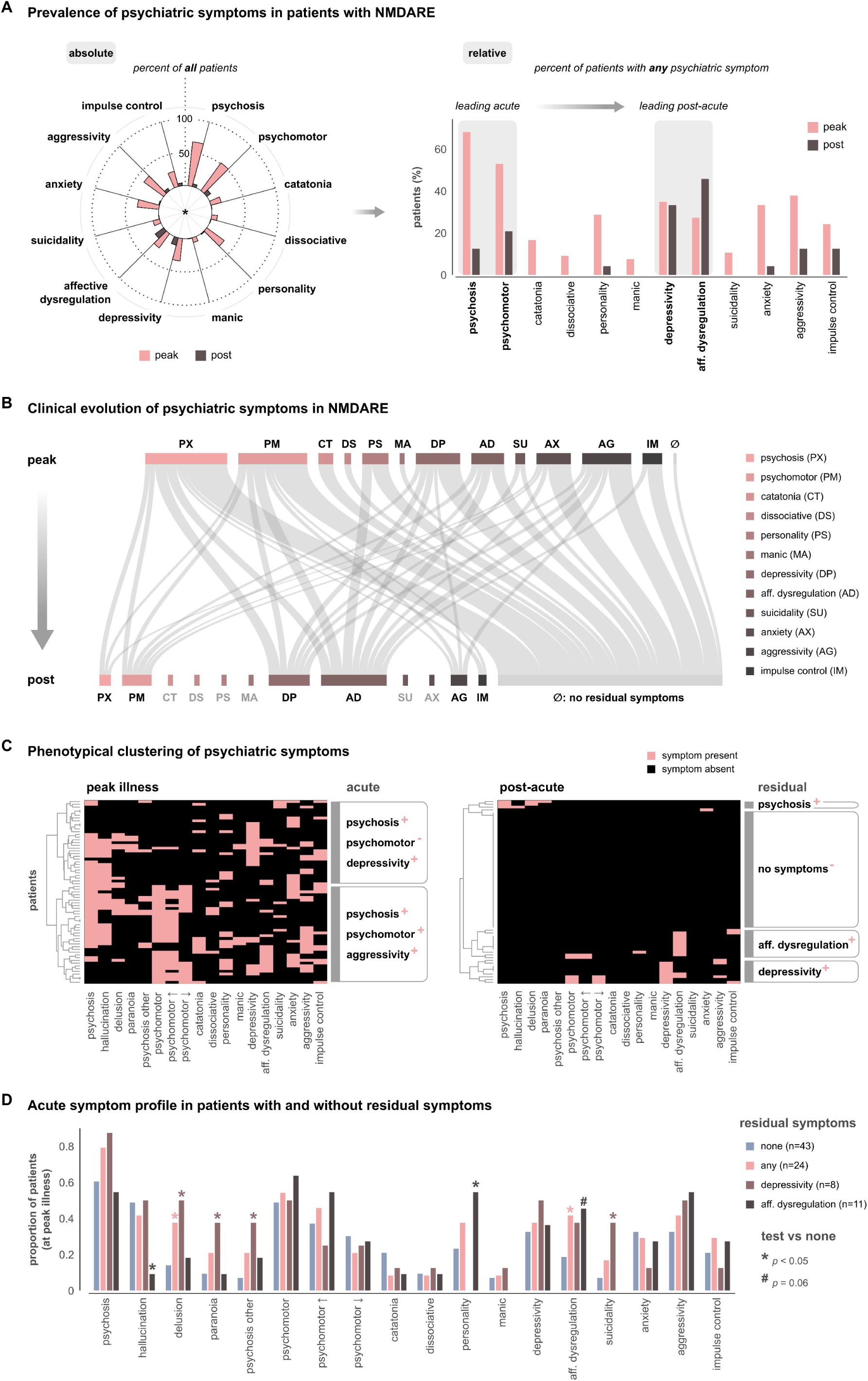
Residual psychiatric symptoms in NMDARE show a phenotypical shift from a ‘schizophrenia-like’ to an affective phenotype. **(A)**, Prevalence of psychiatric symptoms at peak illness and post-acute visit across twelve clinical domains. The polar plot shows the absolute symptom prevalence across all patients (left), and the bar plot visualizes the relative distribution across patients with any psychiatric symptom at each timepoint (right). **(B)**, Sankey plot visualizing the clinical evolution of psychiatric symptoms from peak illness to the post-acute stage. The flow diagram was thresholded to >1 (i.e., links representing only one patient are omitted). **(C)**, Phenotypical clustering of the symptom matrices (patients x dimensions) at peak illness (left) and post-acute visits (right). Clustergrams are based on Euclidean dissimilarity and hierarchical clustering with Ward linkage. **(D)**, Distribution of acute symptoms in patients with and without residual symptoms in the post-acute stage. Abbreviations: **AD**, affective dysregulation, which predominantly included emotional lability (“mood swings”) and to a lesser extent affective flattening; **AG**, aggressivity; **AX**, anxiety-related symptoms; **CT**, catatonia; **DP**, depressivity; **DS**, dissociative symptoms, which predominantly included derealization and depersonalization; **IM**, impaired impulse control; **MA**, manic symptoms; **PM**, psychomotor symptoms; **PS**, personality changes; **PX**, psychosis; **SU**, suicidality, including suicidal behavior, ideation, and clinically reported self-harming tendencies.

However, more than half the patients were completely free of psychiatric symptoms at post-acute follow-up. Therefore, we additionally examined the relative symptom prevalence across patients with *any* psychiatric symptom at each timepoint. This confirmed psychosis and psychomotor symptoms as the leading symptoms in the acute phase and revealed a shift towards depressivity and affective dysregulation as the leading residual symptoms in the post-acute phase (**Fig. 5A**, *right*). Notably, while all other symptoms decreased substantially, the relative prevalence remained stable for depressivity and even increased for affective dysregulation. On the group level, psychiatric symptoms in NMDARE thus showed a phenotypical shift from an acute ‘schizophrenia-like’ phenotype to a residual affective phenotype.

Given these results, we next asked if residual symptoms tend to persist through the acute phase (i.e., they were already present at peak illness) or if they predominantly develop from other symptoms (**Fig. 5B**). Therefore, we analyzed the temporal evolution of these symptoms and observed that (i) patients generally showed improvement across all symptom domains, (ii) some symptoms such as catatonia, dissociative, or manic symptoms resolved completely in all patients, and (iii) residual depressivity and affective dysregulation did not typically persist through the acute phase but rather developed from a wide spectrum of other acute symptoms (**Fig. 5B**, *bottom*).

Consequently, we asked whether these symptom domains tend to coexist in the same patients or rather reflect distinct clinical subgroups (**Fig. 5C**). Phenotypical cluster analysis revealed two larger groups at peak illness: one with low prevalence of psychomotor symptoms and high depressivity, and another with high prevalence of psychomotor symptoms and aggressivity, while psychosis was highly prevalent in both (**Fig. 5C**, *left*). Conversely, in the post-acute stage, symptom clustering suggested four phenotypical groups, where the largest cluster represented patients without any residual psychiatric symptoms, in line with the above analyses and Figure 4C. However, we also observed one small cluster of residual psychosis and two larger clusters characterized by residual depressivity and affective dysregulation, respectively (**Fig. 5C**, *right*). Notably, nearly all these patients showed only one of the latter symptoms, but not both, suggesting that depressivity and affective dysregulation may represent two distinct residual phenotypes.

Therefore, we investigated whether the distribution of acute symptoms differed between patients with residual symptoms and those without (**Fig. 5D**). Indeed, patients with residual depressivity had experienced significantly higher rates of suicidality as well as delusions, paranoia, and ‘other’ psychotic symptoms at peak illness, where the latter predominantly included thought disorders. In contrast, patients with residual affective dysregulation were characterized by significantly lower rates of hallucinations, higher rates of personality changes, and a trend towards higher affective dysregulation in the acute phase.

### Memory performance at MRI

Finally, we focused on patients’ post-acute memory performance. While the analyses in Figure 4C revealed a consistent link between structural brain complexity and memory dysfunction, CASE scoring of memory symptoms is relatively coarse, based on an ordinal scale, and does not differentiate between different memory domains. Therefore, we additionally analyzed patient performance in neuropsychological tests of visuospatial memory (ROCF) and verbal episodic memory (RAVLT) that were prospectively collected at the time of MRI. These test scores are summarized in **Table 2**.

**Table 2.** Memory performance of patients with NMDARE at post-acute visit.

| | No. of patients<br>n (%) | Test score<br>(Mean ± SD) |
| --- | --- | --- |
| **Visuospatial memory** |  |  |
| ROCF: immediate recall | 66 / 70 (94) | 25.9 ± 7.7 |
| ROCF: delayed recall | 66 / 70 (94) | 25.6 ± 7.6 |
| delayed recall: low | 13 / 66 (20) | 13.2 ± 4.9 |
| delayed recall: high | 34 / 66 (52) | 31.3 ± 2.5 |
| **Verbal episodic memory** |  |  |
| RAVLT: sum score | 68 / 70 (97) | 54.4 ± 11.3 |
| RAVLT: interference | 68 / 70 (97) | 10.9 ± 3.9 |
| RAVLT: delayed recall | 68 / 70 (97) | 10.9 ± 3.8 |
| delayed recall: low | 27 / 68 (40) | 7.0 ± 2.5 |
| delayed recall: high | 22 / 68 (33) | 14.7 ± 0.5 |

Abbreviations: No., number; RAVLT, Rey Auditory Verbal Learning Test; ROCF, Rey–Osterrieth complex figure; SD, standard deviation. Interpretation of the delayed recall scores was based on patients' individual percentile ranks in comparison to normative reference values, where low performance was defined as scores <25<sup>th</sup> percentile and high performance as $\geq 75^{\text{th}}$ percentile.

In general, we observed substantial variability in both visuospatial and verbal memory performance. However, age-specific normative interpretation showed that patients were less affected in the domain of visuospatial memory, with only 20% meeting the low-performing criterion compared to their individual normative reference. In contrast, 40% of patients fulfilled the low-performing criterion for verbal memory. Consequently, a significantly greater proportion of patients met the low-performing criterion for verbal than for visuospatial memory (27/68 vs. 13/66, χ^2^=6.40, *p*=0.011), suggesting that residual memory symptoms at MRI are domain-specific, with verbal memory being more affected than visuospatial memory.

Given these findings, we hypothesized that patients with lower post-acute memory performance would show stronger alterations of structural brain complexity, and that this effect would be more pronounced for verbal memory than for visuospatial memory. Indeed, **Figure 6** shows that lower verbal memory performance was associated with higher FD deviation indices, and this was consistently observed for the RAVLT sum score (*r*=-0.33, *p*=0.006, *p_FDR_*=0.008; **Fig. 6A**), interference trial (*r*=-0.41, *p*=4.8*10^-4^, *p_FDR_*=0.002; **Fig. 6B**), and delayed recall (*r*=-0.36, *p*=0.003, *p_FDR_*=0.006; **Fig. 6C**). Moreover, subgroup analysis based on normative references (cf. Table 2) revealed stronger alterations of structural brain complexity in RAVLT low-performers compared to high-performers (*t*=-2.35, *p*=0.023, *d*=-0.68; **Fig. 6D**).

**Figure 6.**
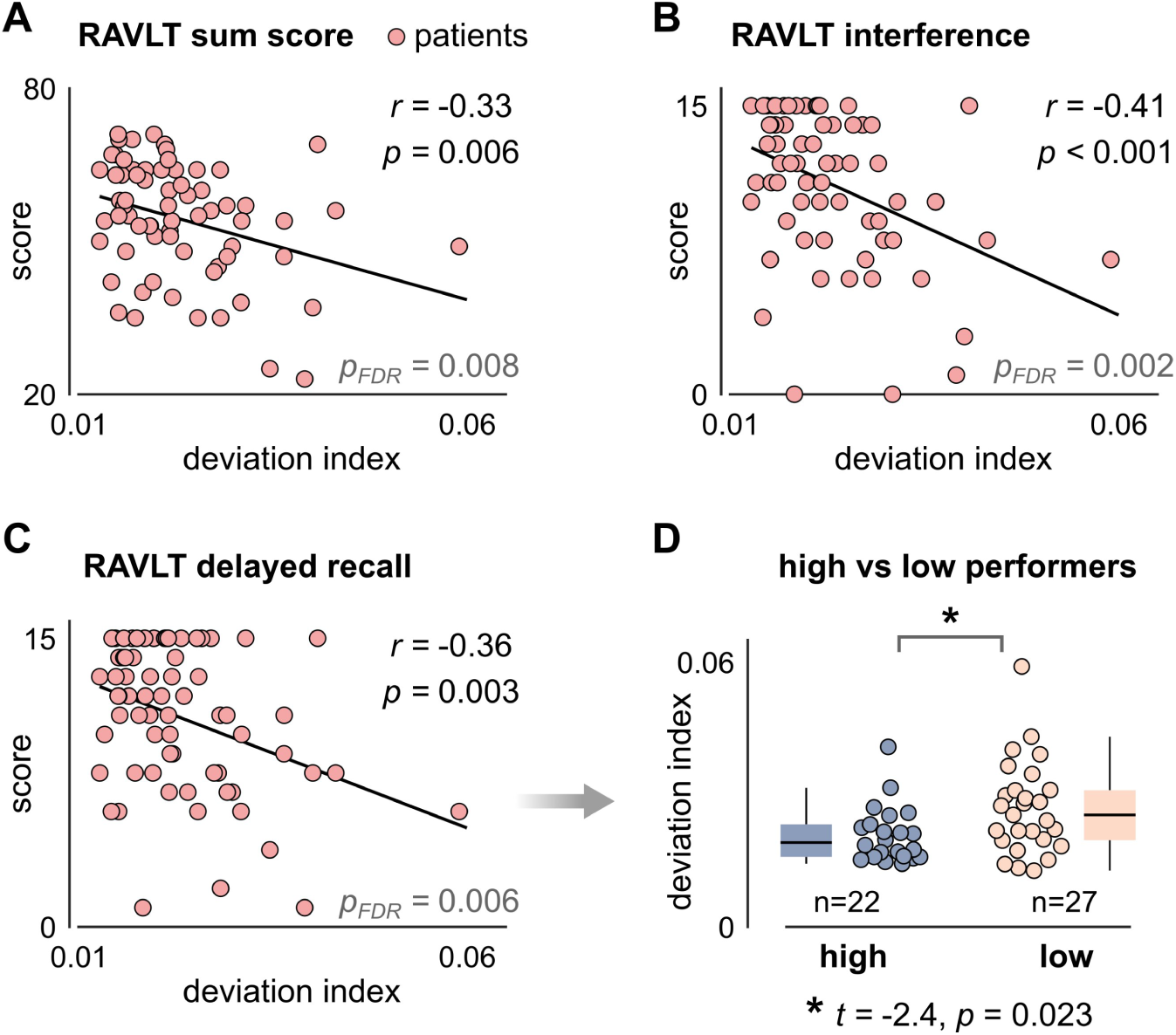
Patients with lower post-acute memory performance show stronger alterations of structural brain complexity. **(A)**, Association between the fractal dimensionality (FD) deviation index (see Fig. 4A) and patients’ verbal episodic memory performance as assessed by the Rey Auditory Verbal Learning Test (RAVLT) sum score. **(B)**, Association between the FD deviation index and RAVLT post-interference performance. **(C)**, Association between the FD deviation index and RAVLT delayed recall scores. **(D)**, Subgroup analysis comparing patients who met the high-performing criterion (≥75th percentile of their age-specific normative reference) and those who met the low-performing criterion (<25th percentile) in the delayed recall scores (see Table 2).

Additionally, we observed similar effects for the domain of visuospatial memory, albeit with slightly lower effect sizes: Here, stronger alterations of structural brain complexity were significantly associated with lower ROCF scores, both for immediate recall (*r*=-0.30, *p*=0.014, *p_FDR_*=0.014) and delayed recall performance (*r*=-0.35, *p*=0.004, *p_FDR_*=0.006). Moreover, ROCF low-performers showed stronger FD alterations than high-performers (*t*=-2.03, *p*=0.048, *d*=-0.66).

Notably, these effects were not driven by age differences between high- and low-performers, neither for verbal memory (*t*=-0.03, *p*=0.98), nor visuospatial memory (*t*=0.17, *p*=0.87). Similarly, we observed no sex differences between low- and high-performers, although there was a trend towards a higher proportion of males in the low-performing subgroup of verbal memory (ROCF: χ^2^=0.02, *p*=0.90; RAVLT: χ^2^=2.29, *p*=0.13).

## Discussion

This study combined clinical analyses, neuropsychological assessments, and advanced neuroimaging techniques to show that memory dysfunction and psychiatric outcomes are linked to altered structural brain complexity in post-acute NMDARE. Specifically, we leveraged the recent framework of fractal dimensionality analysis and observed that (i) patients showed systematic reductions of structural brain complexity compared to age- and sex-matched healthy individuals; (ii) these reductions followed a characteristic spatial pattern including the hippocampus and a fronto-cingulo-temporal cluster in both cerebral cortex and white matter; (iii) two-thirds of patients experienced residual symptoms despite immunotherapy and good overall clinical recovery; (iv) the most common residual symptoms were memory dysfunction and psychiatric symptoms; (v) psychiatric symptoms showed a phenotypical shift from a ‘schizophrenia-like’ phenotype at peak illness to an affective phenotype in the post-acute stage; and (vi) patients with residual symptoms displayed stronger alterations of structural brain complexity than those without. Collectively, these findings advance our understanding of NMDARE along four key directions.

### Structural complexity reductions

First, we show that structural brain changes in post-acute NMDARE follow a highly specific spatial pattern, uncovered through fractal dimensionality (FD) analysis of anatomical MRI data. Therein, FD quantifies the geometric complexity of a brain structure, which is mathematically distinct from its size and where higher values reflect more irregular shapes.^16,17^ Here we found that NMDARE is characterized by systematic reductions of structural brain complexity, which cluster in previously implicated brain regions but also identify several new disease correlates.

Overall, the strongest FD reductions were observed in the hippocampus bilaterally. These effects align well with previous neuroimaging studies reporting (i) volume reductions^33,34^, (ii) impaired microstructural integrity^33,35^, (iii) surface deformations^34^, and disrupted functional connectivity of the hippocampus^36–39^. Our results thus clearly corroborate the hippocampus as a major disease target and additionally show that hippocampal changes in NMDARE include long-term shape alterations besides volume loss, microstructural impairments, and connectivity alterations.

In addition, we identified a fronto-cingulo-temporal cluster of FD reductions, which involved both cortical GM and cerebral WM and mapped onto high-order functional systems. Regarding the WM changes, our results align well with previous neuroimaging studies reporting widespread WM damage in NMDARE^35,36,40,41^. Of particular interest, one study found that non-recovered patients showed extensive damage of late-myelinating superficial WM, a particularly vulnerable tissue compartment^42^. Notably, these changes clustered in frontal and temporal lobes, closely matching the spatial pattern of WM changes observed here. Additionally, however, our results implicate cingulate WM, where prior studies have found evidence of impaired tissue integrity^36,41^. Our results thus align well with previously reported patterns of WM damage. However, this convergence is remarkable given that prior studies used a fundamentally different imaging modality –diffusion tensor imaging– commonly used to assess WM *micro*structure. In contrast, our results reveal a systematic pattern of *macro*structural WM changes in NMDARE, rooted in altered WM morphology.

Moreover, we observed a similar fronto-cingulo-temporal cluster of FD reductions in the cerebral cortex. These alterations are particularly interesting as cortex morphology in NMDARE has received comparatively less attention and several prior studies have not detected any cortical changes in standard volumetry^36,43^. However, in two more recent studies that did find volume reductions in the cortex, changes were also most prominent in the frontal and temporal lobes^44,45^ where the density of NMDA receptors is thought to be particularly high^46^. While cortical volume changes in NMDARE may thus be more variable (or transient), structural complexity analysis clearly identified long-term frontal and temporal changes in the cerebral cortex. Importantly, however, the strongest cortical effects were again observed in the cingulate. Although we are unaware of any prior studies implicating the morphology of the cingulate cortex in NMDARE, one recent study reported functional changes in the cingulate that were related to cognitive impairment^47^. As such, the strong effects in cingulate GM and WM observed here are particularly compelling, given that (i) the cingulate is commonly viewed as a hub region for both memory function and affective processing^48^ –which represented the two leading domains of residual symptoms in our cohort– and (ii) microstructural changes in the cingulate have been reported in both schizophrenia and depression^49^, aligning well with our results on psychiatric outcomes.

### Psychiatric outcomes

Second, we show that changes in structural brain complexity are significantly stronger in patients with residual psychiatric symptoms than in those without. Notably, psychiatric manifestations represented the leading CASE domain at peak illness and the second most prevalent residual symptoms in the post-acute stage, such that we characterized the temporal evolution of these symptoms in detail. Therein, the symptom domains identified in our cohort aligned remarkably well with those reported in a large-scale analysis of NMDARE-related psychopathology, which found that core aspects of mood disorders and psychosis consistently coexist in these patients^50^.

Importantly, our study confirms this mixed mood-psychosis syndrome and additionally shows that psychiatric symptoms dynamically evolve from a ‘schizophrenia-like’ phenotype at peak illness to an affective phenotype in patients with residual psychiatric symptoms.

In general, the transdiagnostic relationship between NMDARE and schizophrenia has long been recognized, including overlapping clinical features and disruptions in glutamatergic signaling as a potential shared pathomechanism^51,52^. In this context, one key outcome study found that cognitive and psychiatric symptoms in the early post-acute stage of NMDARE indeed resemble those of stabilized schizophrenia.^11^ However, recovery in that cohort was not linear, with the most pronounced improvements occurring between 4 and 10 months after hospital discharge. Given the somewhat longer disease duration in our study (on average ∼2 years since onset), this suggests that the shift towards residual affective symptoms may reflect ongoing (and potentially modifiable) disease processes in patients who continue to experience psychiatric symptoms in the post-acute stage.

Strikingly, both low mood and affective dysregulation featured prominently in a recent study on patient-reported outcomes of NMDARE, where subjective depressive symptoms were furthermore associated to lower self-reported quality-of-life.^13^ These self-reports not only strengthen the clinical relevance of our findings but also highlight the need for prognostic markers of long-term psychiatric outcomes.

In this context, our results suggest that the presence of certain symptoms at peak illness may indicate an elevated risk of residual psychiatric symptoms in the post-acute stage. In particular, we found that residual depressivity in the post-acute stage was associated with the presence of delusions, paranoia, thought disorders, and self-harming tendencies in the acute phase. Furthermore, patients with affective dysregulation in the post-acute stage were characterized by disproportionately higher rates of personality changes and, interestingly, a relative lack of hallucinations in the acute stage. Although dedicated studies will be necessary to evaluate such clinical predictors systematically, these results further support the phenotypical shift in psychiatric manifestations of NMDARE and show that clinical phenotyping represents a promising avenue to identify at-risk profiles and derive more personalized prognostic markers.

### Memory dysfunction

Third, we found that patients with residual memory dysfunction show stronger alterations of structural brain complexity than those without. Notably, memory dysfunction was highly prevalent at peak illness and represented the leading CASE domain of residual symptoms in the post-acute stage. These results closely match previous evidence showing that long-term memory dysfunction is common in NMDARE^9,12,53,54^ and represents one of the most important patient-reported concerns^13^.

Additionally, our study found that patients with stronger brain alterations performed worse on prospective memory tests, where verbal episodic memory was comparatively more affected than visuospatial memory. In this context, it has been argued that some cognitive domains such as episodic memory may be less likely to recover than others^54^, and the relatively stronger impact on verbal episodic memory was similarly observed in a recent related study^9^. Notably, however, fractal analysis sensitively detected stronger brain changes in more affected patients across both the broader CASE score of memory dysfunction and the domain-specific memory assessments at MRI, yielding several implications for future research and post-acute monitoring.

### Clinical implications

Fourth, our study supports the view that current treatment strategies do not sufficiently prevent long-term sequelae in many patients with NMDARE. Consequently, the disease has been characterized as only partially responsive to immunotherapy^8^, highlighting the need for better prognostic markers and improved post-acute monitoring.

In this regard, our study offers FD as a promising new imaging marker to track clinically relevant brain changes. Importantly, FD does not require specialized MRI sequences but can be derived from standard 3D-T1-weighted images, facilitating its use in future studies and, potentially, clinical practice. This advantage is met by recent methodological advances that leverage artificial intelligence to convert clinical MRIs with lower resolution into research-grade high-resolution scans^55^.

Additionally, the normative modelling framework applied here offers a principled way to derive more personalized outcome markers, as it explicitly allows for patient-specific patterns of brain alterations. Besides the current study, this approach has shown high sensitivity in a recent study on neurodevelopment^17^ and a related application in anti-LGI1 encephalitis^29^. Relating a patient’s individual scan to a demographically matched healthy reference thus represents a promising avenue to track disease-related brain changes and treatment effects in future studies.

Despite these promising implications, limitations of our study include the cross-sectional design, which precludes inferences on when during the disease course the observed brain alterations develop. Similarly, patients were included upon referral to our reference center, such that time intervals since the acute disease phase varied. Although this variability did not have a significant impact on residual symptoms in our study, longitudinal assessments will be necessary to understand symptom trajectories in more detail and to characterize the temporal dynamics of the observed phenotypical shift in psychiatric symptoms.

In sum, our results carry immediate implications for understanding the neuropathology, clinical evolution, and outcomes of NMDARE and highlight FD as a promising imaging marker to capture patient-specific patterns of brain alterations using routine MRI data.

## Notes

### Competing Interest Statement

The authors have declared no competing interest.

